# Binocular Benefit Following Monocular Subretinal AAV Injection in a Mouse Model of Autosomal Dominant Retinitis Pigmentosa (adRP)

**DOI:** 10.1101/2021.07.22.453413

**Authors:** Chulbul M Ahmed, Michael T Massengill, Cristhian J Ildefonso, Ping Zhu, Hong Li, Anil P. Patel, Alfred S Lewin

## Abstract

Autosomal dominant retinitis pigmentosa (adRP) is frequently caused by mutations in *RHO*, the gene for rhodopsin. In previous experiments in dogs with the T4R mutation in *RHO*, an AAV2/5 vector expressing both an shRNA directed to human and dog *RHO* mRNA and an shRNA-resistant human *RHO* cDNA (AAV-*RHO820-shRNA820*) prevented retinal degeneration for more than 8 months following injection. To confirm that this same vector could protect the retinas of a different species and bearing a different *RHO* mutation, we injected mice transgenic for human P23H *RHO* at postnatal day 30 in one eye. For nine months, we monitored their retinal structure using spectral-domain optical coherence tomography (SD-OCT) and retinal function using electroretinography (ERG). We compared these to P23H *RHO* transgenic mice injected with AAV-GFP. Though retinas continued to thin over time, compared to control injected eyes, AAV-RHO820-shRNA820 slowed the loss of photoreceptor cells and decreased ERG amplitudes in AAV-*RHO820-shRNA820* eyes during the nine-month study period. Unexpectedly, we also observed preservation of retinal structure and function in the untreated contralateral eyes of AAV-*RHO820-shRNA820* treated mice. PCR analysis and western blots provided evidence that a low amount of vector from injected eyes was present in uninjected eyes.

## Introduction

Retinitis pigmentosa is the name for a collection of inherited diseases leading to vision loss caused by the death of photoreceptor cells of the retina^1^. The disease gets its name from the deposition of pigment in the retina that can be seen in a routine fundus examination. While phenotype is variable, even among members of the same family, symptoms usually begin with night blindness and loss of peripheral vision. The disease may stop at that point, but in many people, loss of retinal function is progressive, leading to tunnel vision or to near total loss of light sensitivity. Although inheritance may be X-linked or recessive, about one-third of retinitis pigmentosa cases are inherited dominantly. Mutations in the gene for rhodopsin, *RHO*, are responsible for approximately one-quarter of the autosomal dominant retinitis pigmentosa (adRP) cases.

Previously published research described a combination therapy for adRP using mouse models of this disease that were transgenic for a mutant version of human *RHO* containing the proline to histidine mutation at residue 23 (P23H), the most prevalent mutation among adRP patients in North America^2^. In our approach, a single adeno associated virus (AAV) vector expressed both an shRNA targeted to human *RHO* and a *RHO* gene containing nucleotide changes that prevented binding of the siRNA product without changing the amino acid sequence of *RHO*. In our experiment, retinal degeneration halted by two months post-treatment, and the preservation of retinal structure and function lasted for nine months when the experiment ended. Another group used two AAV vectors to achieve retinal protection in mice with a different adRP mutation^3^. More recently, Tsai and colleagues have used CRISPR-Cas9 to disrupt the mouse P23H *Rho* gene in a knock-in model of adRP, replacing the missing rhodopsin via a human *RHO* cDNA that is resistant to the guide RNAs used for ablation of the mouse gene^4^.

Because of our success in the mouse model, we joined with investigators at the University of Pennsylvania who had carefully characterized dogs with the heterozygous T4R mutation in *RHO*^5^. This change alters a glycosylation site in rhodopsin and leads to slow progressive retinal degeneration in ambient lighting. When the eyes of T4R *RHO* dogs are briefly exposed to intense light, however, retinal degeneration ensues immediately, leading to total loss of photoreceptor cells in the illuminated area^6^. We previously screened a series of siRNAs and ribozymes targeting human and canine *RHO* mRNA to identify shRNA820 as particularly effective in diminishing the level of canine rhodopsin without affecting photoreceptor viability following subretinal delivery by AAV2/5.

The siRNA derived from this shRNA binds the mRNA surrounding nucleotide 820 in the human *RHO* mRNA, a site unaffected by adRP mutations, and an equivalent site in the canine *RHO* mRNA. A combination vector expressing this shRNA and a resistant human *RHO* cDNA were injected subretinally in *T4R RHO* dogs under infrared illumination, and the injected area was protected from retinal degeneration following subsequent multiple rounds of illumination^7^.

The effectiveness of the combination vector, designated scAAV2/5-hOP-*RHO820*-H1- *shRNA820* (shortened here to AAV-RHO820-shRNA820), in the canine model of adRP suggested that it may help treat human adRP caused by *RHO* mutations, since the human *RHO* promoter and transgene are used in this vector, and the shRNA targets human *RHO* mRNA. Before proceeding, we wanted to verify that this approach was mutation independent, that is, that the vector could treat adRP arising from a different *RHO* mutation. We also wanted to confirm that it could treat adRP in an organism other than dogs. For that reason, we have tested it in the P23H *RHO* transgenic mouse model of adRP. In these experiments, we injected only one eye of test mice, comparing scAAV2/5-hOP-*RHO820*-H1-*shRNA820* injected eyes with AAV-GFP infected eyes of a separate group of mice. We found that the combination therapy led to improved retinal structure and function compared to the control eyes for the duration of the experiment (9 months). Unlike our previous RNA replacement therapy in adRP mice and dogs, we observed comparable protection in the uninjected, contralateral eyes of mice treated in one eye with AAV-RHO820-shRNA820, even though we detected very low levels of RHO820 and AAV vector in those eyes.

## Results

### ShRNA820 degrades both wild type and P23H human RHO mRNA

Our previously published transfection experiments demonstrated that shRNA820 leads to cleavage of both wild type and P23H *RHO* mRNA. ^7^ There is a one nucleotide change between human and mouse rhodopsin mRNA at position 5 of the shRNA target site (**Fig. 1A**). This change was sufficient to prevent cleavage directed by shRNA 820, but when we tested a perfectly matched mouse version of shRNA 820, we observed an efficient blockade of RHO-GFP expression (**Fig. 1B**). Based intensity of the monomer rhodopsin band, the mouse shRNA820 led to a 78% knockdown of *Rho* mRNA, whilst human shRNA820 caused no significant change (**Fig. 1C**).

**Fig. 1.**
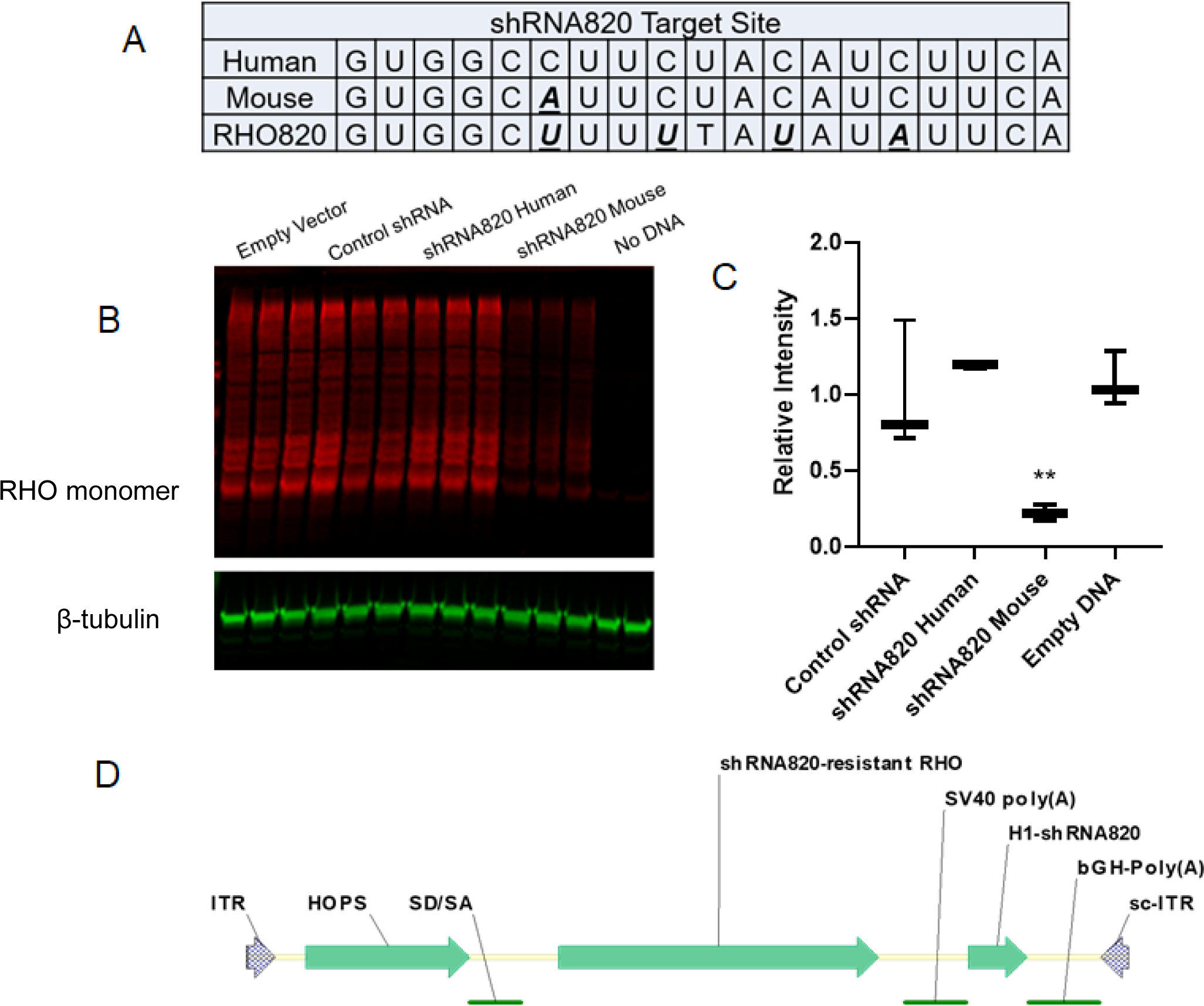
One mismatch with Mus Rho abolishes the activity of shRNA820. **A**. One mismatch occurs between mRNAs of human RHO and mouse Rho. The shRNA resistant RHO820 mRNA contains four changes relative to wild type human RHO mRNA. The sequence of WT-huRHO in the CMV-WT-huRHO-tGFP plasmid was exchanged for the coding sequence of musRho from and a mouse version of the H1-shRNA820 cassette was generated and cloned in the SalI site of an AAV vector (sc-rAAV2-MOPS-reverseGFP). 500ng of either the CMV-Rho-tGFP as well as 1µg of one of the shRNA plasmids, including control, were co-transfected in HEK293T cells and incubated for 48 hours before assaying for knockdown. An sc-rAAV2 plasmid without an shRNA (empty DNA) was also employed as a control. **B.** Protein was extracted from the cells and analyzed by western blotting, probing for TurboGFP (red) and β-tubulin (green). **C.** We measured band intensities corresponding to the predicted monomeric form of huRHO-tGFP (65kDa) or all aggregated forms of huRHO-tGFP and corrected for loading by dividing by the band intensity of tubulin of the same well. The relative intensity was calculated by dividing the corrected band intensity of a given well by that of the control shRNA condition. The bar graph and error bars represent the mean ± SEM of three biological replicates. We used one-way ANOVA followed by multiple comparison tests to determine statistical significance. ** = p<0.01. **D.** Map of AAV-RHO820-shRNA820: ITR= inverted terminal repeat; sc-ITR=internally deleted ITR; HOPS=human opsin proximal promoter; SD/SA= splice donor and splice acceptor from SV40; H1-shRNA820=shRNA820 cassette driven by the human H1 promoter.

### AAV2/5-RHO820-shRNA820 protects the retina of P23H *RHO* transgenic mice

We constructed a combination vector containing shRNA820 driven by the human H1 promoter and an siRNA820- resistant human *RHO* cDNA driven by the human opsin proximal promoter (**Fig. 1D**). The resistant gene, which we call RHO820, has four silent base pair differences from human *RHO* mRNA (Fig. 1A). This combination vector (AAV-RHO820-shRNA820) has several significant differences from the similar AAV vector we tested previously in P23H *RHO* transgenic mice^2^. The shRNA and, therefore, the resistance mutations are different. The transgene is derived from human rhodopsin, whereas our previous experiments used the mouse *Rho* cDNA. The promoter for rhodopsin expression is based on the proximal promoter sequences of the human *RHO* gene, while previously we used the mouse *Rho* proximal promoter. We previously used a single-stranded AAV2/5 vector, but our current vector is self-complementary due to a deletion in the 3’ ITR sequence^8^.

We injected 1µl (5 x 10^8^ vector genomes) of this vector subretinally in the right eyes of one month old P23H *RHO* transgenic mice (N=20). Left eyes were not injected. As a control for the impact of AAV treatment, we injected the same volume and titer of the AAV2/5-GFP vector, in which the GFP gene was under the control of the CBA (CMV enhancer-chicken β-actin) promoter, in the right eyes of a separate cohort of one month old mice (N=15). We viewed the AAV-GFP injected eyes by fluorescence fundoscopy one month post injection to validate our injection technique (**Supplemental Fig. 1**). We observed more than 70% transduction in the observed fields (central 30% of the retina) in the GFP injected eyes. No GFP fluorescence was detected in the contralateral eye. Injection of AAV-RHO820-shRNA820 led to a 20% reduction of total human RHO mRNA in the injected eyes relative to the uninjected eyes of the same mice (p<0.001), based on qRT-PCR using human *RHO* specific primers (hRHO-F and hRHO-R in Table 1) one month post-injection (**Fig. 2A**). This result indicates that the amount of rhodopsin mRNA produced from the *RHO*820 cDNA delivered by the vector did not fully replace the amount of P23H *RHO* mRNA depleted by the siRNA820 (the guide strand of shRNA820) in these mice. In addition, the level of *RHO* mRNA in the *uninjected* eyes was significantly less than that detected in the eyes of naïve (untreated) P23H *RHO* transgenic mice of the same age (p<0.01). This result suggests that knockdown of P23H *RHO* RNA also occurred in the uninjected eyes of AAV-RHO820-shRNA820 treated mice. However, we did not detect a difference in total rhodopsin protein (mouse plus human) between injected and uninjected eyes by western blot performed at the same time (**Fig. 2B and 2C**).

**Fig. 2.**
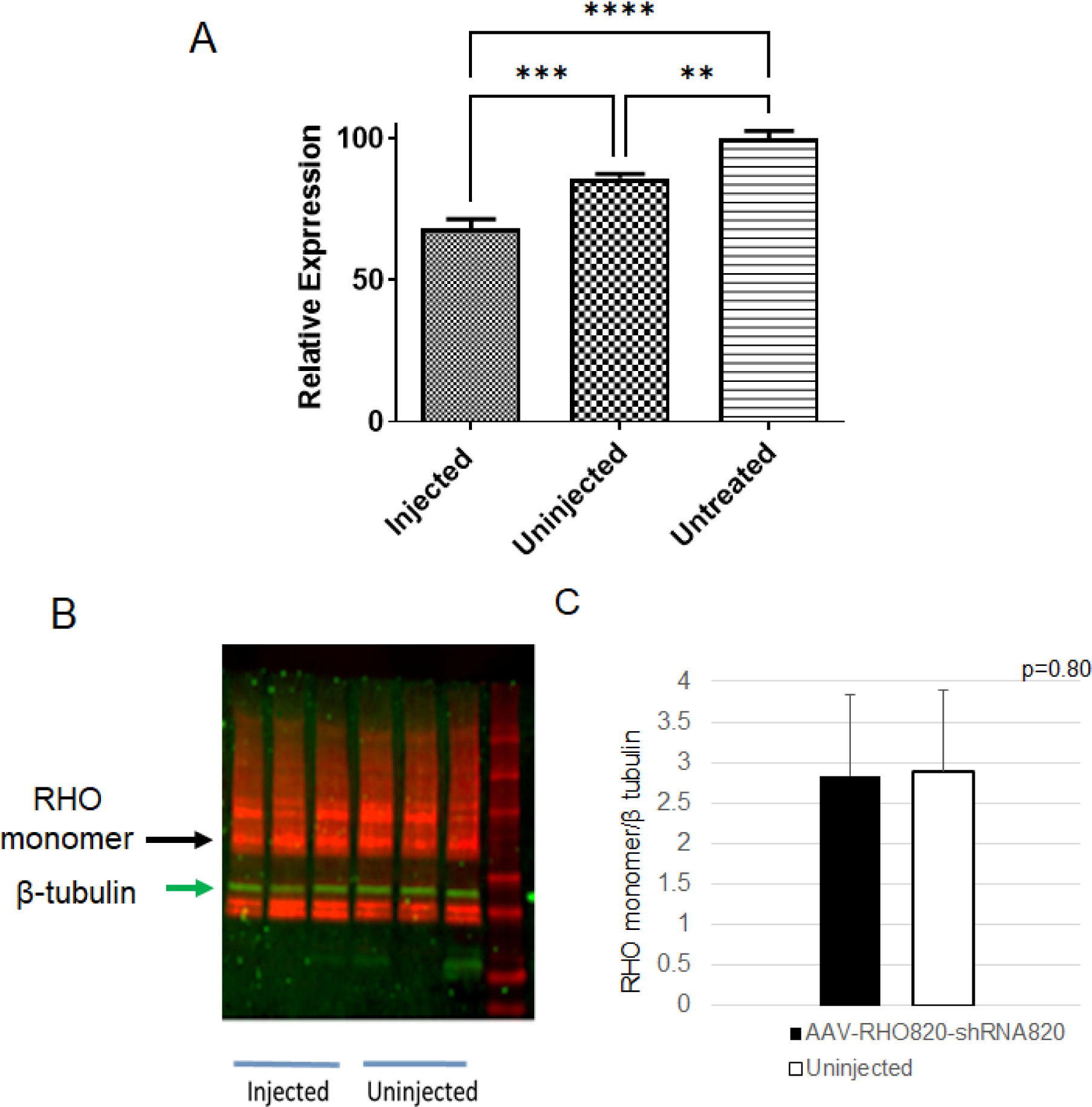
Subretinal injection of AAV-RHO820-shRNA820 leads to the knockdown of human rhodopsin mRNA. **A**. We injected mice unilaterally with AAV-RHO820-shRNA820 and extracted RNA from injected and uninjected eyes, as well as from P23H *RHO* transgenic mice that had not been injected with the virus. Human RHO mRNA was amplified using the primers hRHO-F and hRHO-R listed in Table 1 (n=3), normalizing the transcript levels to β-actin. Error bars indicate standard deviation. **B**. We dissected retinas treated and untreated eyes of AAV-RHO820- shRNA820 injected mice and separated the proteins on an SDS-polyacrylamide gel, which was subsequently transferred to a PVDF membrane and probed with a mouse monoclonal antibody to rhodopsin (Abcam, ab98887) and a monoclonal antibody to β-tubulin. Antigen-antibody complexes were detected with infrared dye-labeled anti-mouse IgG. **C.** We normalized the intensities of the rhodopsin monomer band to that of α-tubulin, and found no differences between injected and uninjected eyes. (We infer that the bands below tubulin, are immunoglobin heavy chains detected by the anti-mouse secondary antibody.)

**Table 1.**
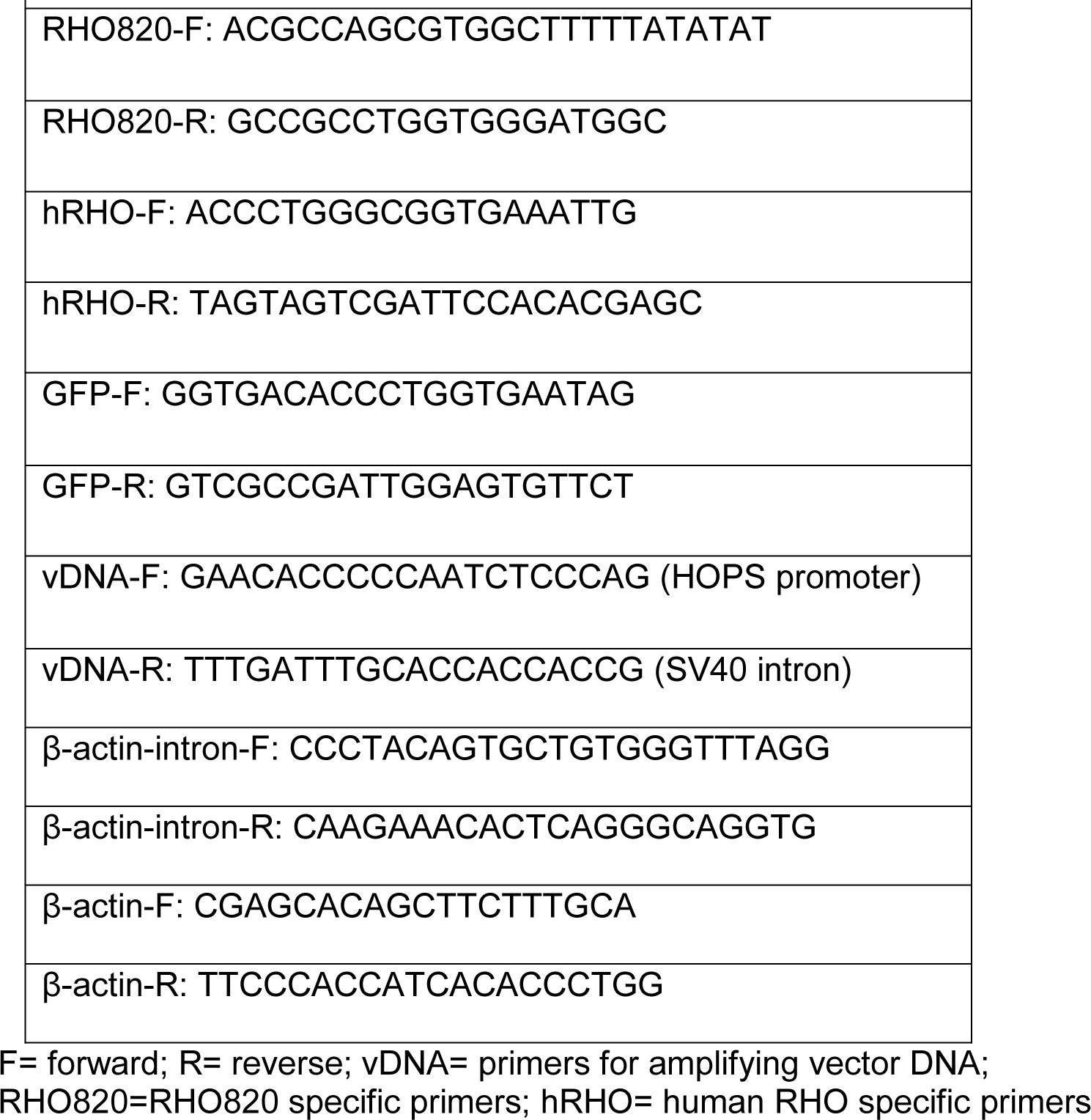
Nucleotide sequences of PCR primers used for qPCR

We measured the integrity of the retinas using spectral domain optical coherence tomography (SD-OCT), a reflectance technique used clinically to assess retinal and optic nerve disease. Images from pre-injected mice and from the first nine months post-injection are shown in **Fig. 3A**. At each time point post-injection, the thickness of the entire retina and of the outer nuclear layer, representing the cell bodies of photoreceptors, was increased in the eyes treated with AAV- RHO820-shRNA820 as compared with eyes injected with AAV-GFP. In addition, the laminar structure of the outer retina was compromised in the control-treated mice, as early as one month post-injection (**Supplemental Fig. 2**). When we quantified the ONL measurements over a nine-month time course (**Fig. 3B**), it was apparent that treatment with a combination vector led to improved integrity of the retina at each interval. By one month post-injection (two months of age) the ONL thickness in control-treated eyes had decreased by 40%, and by three months, half of the photoreceptors had degenerated based on ONL thickness. By nine months, the ONL was barely measurable in control eyes. In contrast, eyes treated with AAV-RHO820-shRNA820 retained half of the starting ONL thickness at nine months post-injection (10 months of age). After the nine-month time course, mice were humanely euthanized and the eyes of some mice were prepared for light microscopy by fixation followed by paraffin embedding. Hematoxylin-eosin staining of sections from these retinas confirmed our conclusions based on SD-OCT. The outer nuclear layers of AAV- RHO820-shRNA820 treated eyes appeared thicker than those of AAV-GFP treated eyes, and the outer segment lengths appeared longer than AAV-GFP injected eyes (**Fig. 3C**).

**Fig. 3.**
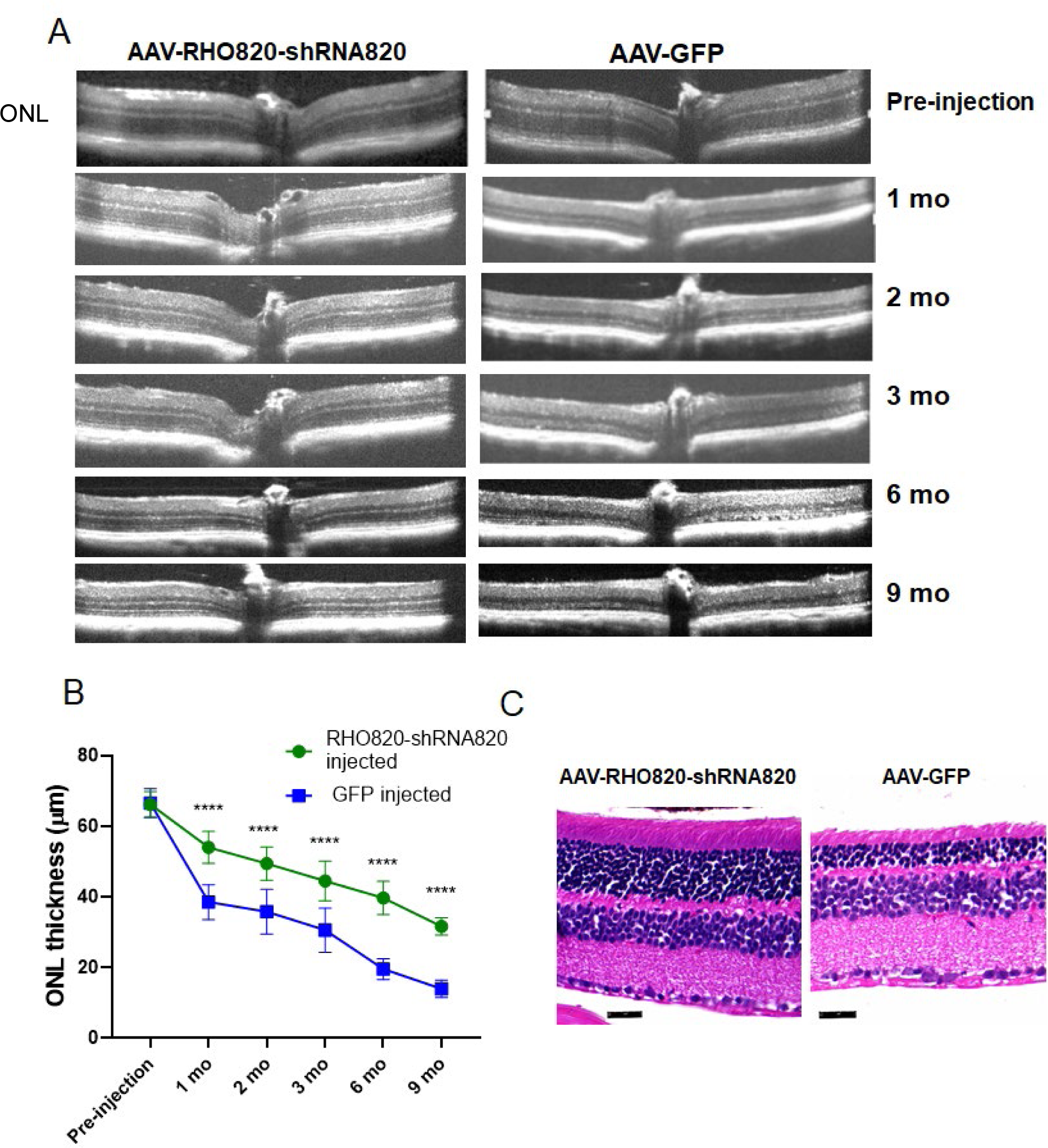
Treatment with the RNA replacement vector preserves outer retinal integrity. **A**. SD-OCT B- scans of eyes treated with AAV-RHO820-shRNA820 or AAV-GFP pre-injection and 1, 2, 3, 6 and 9 months after subretinal injection of one month old mice. **B**. Outer nuclear layer (ONL) thickness was measured at four locations equidistant from the optic nerve head and averaged for each eye. The data points are the averages of 10-15 pairs of eyes; ****=p<0.0001 (Supplemental Table 1). **C**. After the final assessment at nine months post-injection, we fixed the eyes of some mice in paraformaldehyde, embedded them in paraffin and prepared them for staining with hematoxylin and eosin. Representative images of AAV-RHO820-shRNA820 and AAV-GFP injected eyes are shown. The scale bar is 25 micrometers.

### AAV-RHO820-shRNA820 improves the ERG b-wave response of P23H *RHO* transgenic mice

As expected from the loss of ONL thickness, the electroretinogram (ERG) a-wave response, reflecting the function of photoreceptor cells, declined substantially in AAV-GFP treated eyes of P23H transgenic mice. By one month post-injection the a-wave amplitudes decreased by 74%, and they continued to decline until three months post-injection, after which they leveled off at 15% of the pre-injection value (**Fig. 4A**). Treatment with AAV-RHO820-shRNA820 led to increased a-wave amplitudes at all time points, but these increases did not reach statistical significance except at one month and three months. In contrast, the ERG b-wave amplitudes, measuring the responses of secondary neurons in the retina, were significantly higher at all post-injection time points in AAV- RHO820-shRNA820 treated eyes relative to control eyes (**Fig. 4B**). Similarly, there was a statistically significant increase in the light-adapted (photopic) b-wave response from one- to nine-months post-injection in treated eyes, though the difference at six months was not significant (**Fig. 4C**). Overall, the functional preservation afforded by treatment with AAV-RHO820-shRNA820 was less substantial than the structural preservation detected by SD-OCT (**Fig. 3**).

**Fig. 4.**
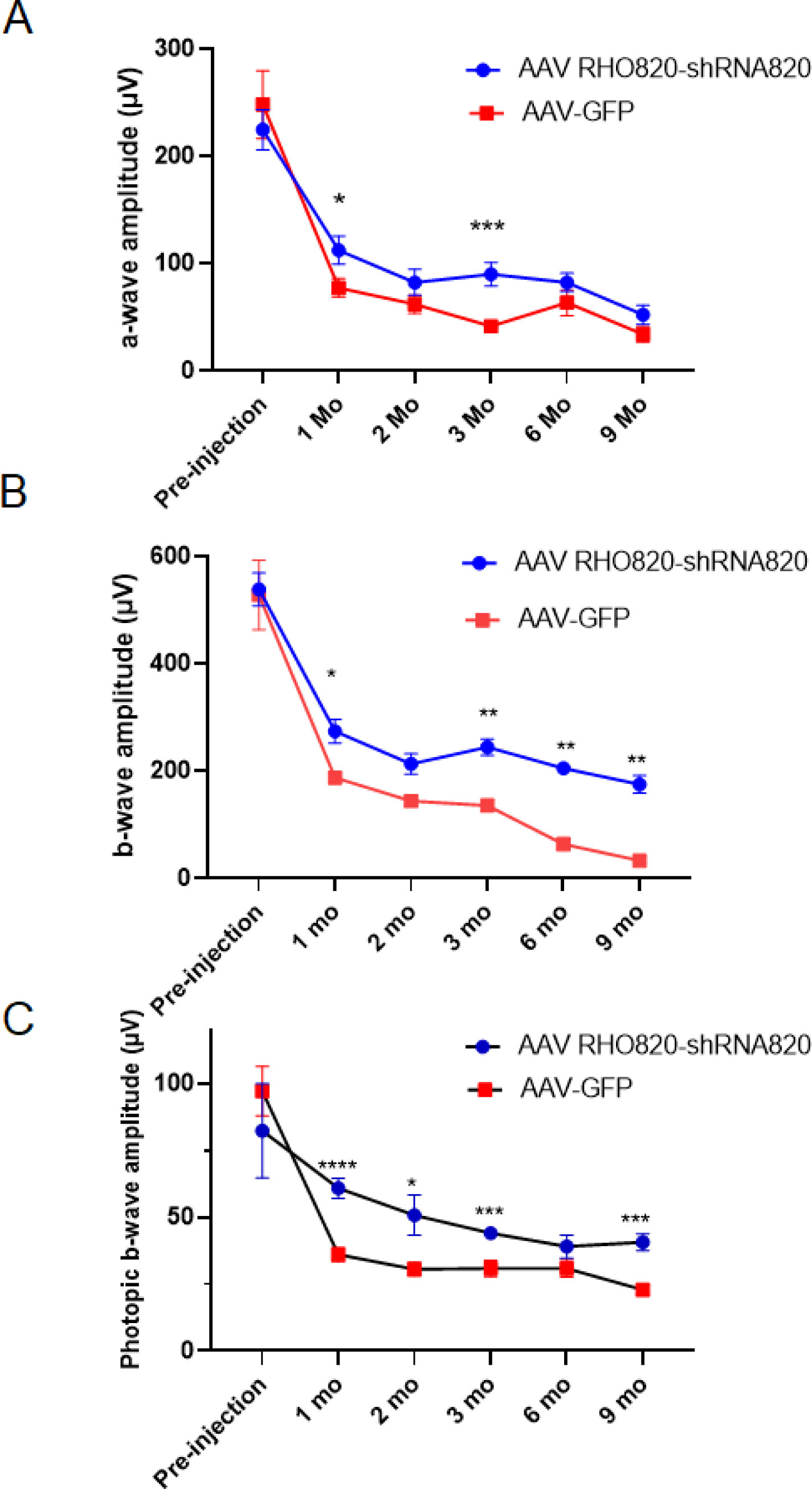
Treatment with AAV-RHO820-shRNA820 protects retinal function. Dark-adapted (scotopic) and light-adapted (photopic) electroretinograms were measured in eyes treated with AAV-RHO820- shRNA820 or with AAV-GFP (n= 14 for pre-injection and months 1-6; n=5 for month 9). The averages of a-wave and b-wave amplitude are plotted. **A**. Scotopic a-wave amplitudes; **B**. Scoptopic b-wave amplitudes; **C**. Photopic b-wave amplitudes. Eyes treated with AAV-RHO820- shRNA820 showed consistently higher amplitudes than eyes injected with AAV-GFP. We used two- way ANOVA followed by multiple comparison tests to determine statistical significance. Differences were statistically significant between AAV-RHO820-shRNA820 and AAV-GFP treated mice at 1, 3, and 6 months for scotopic b-wave amplitudes and at 3 months for scotopic a-wave amplitudes. For photopic amplitudes, differences were significant at months 1-3. .*=p<0.05;** = p<0.01;***=<0.001.

### Protection of contralateral eyes

Since we injected only the right eyes of mice with AAV-RHO820-shRNA820 or AAV-GFP, we were able to measure the impact of treatment on the untreated contralateral eyes of the same mice. We observed the same structural benefit of treatment in the uninjected left eyes as we did in the AAV-RHO820-shRNA820 treated eyes. In contrast, injection of AAV-GFP conferred no benefit to contralateral eyes of the same mice. Protection of the contralateral eyes was unexpected based on our previous results^2, 7, 9^, and those of other testing gene therapy in mouse models of RP^3, 10–12^. However, protection of contralateral eyes is consistent with the knockdown of human *RHO* mRNA in the uninjected contralateral eyes (**Fig. 2A**) The impact of treatment on retinal integrity is illustrated in **Fig.5.** In AAV-RHO820-shRNA820 treated mice, SD-OCT B-scans revealed similar retinal and ONL dimensions in both injected and uninjected eyes (**Fig. 5A**). ONL measurements showed that relative protection of the uninjected eyes persisted for the entire nine-month time course (**Fig. 5B**). Two-way Analysis of Variance indicated no significant difference between AAV-RHO820-shRNA820 injected and uninjected (contralateral) eyes of the same mice (**Supplemental Table 1**). In contrast, AAV-GFP treatment conferred no protection of the contralateral eyes (**Fig. 5B, Supplementary Fig. 3**). Histological analysis performed at nine months post-injection, confirmed the SD-OCT findings: ONL thickness and outer segment lengths appeared shorter in both the AAV-GFP treated eye and untreated eyes compared to either eye of a mouse treated with AAV-RHO820-shRNA820 (**Supplemental Fig. 4**)

**Fig. 5.**
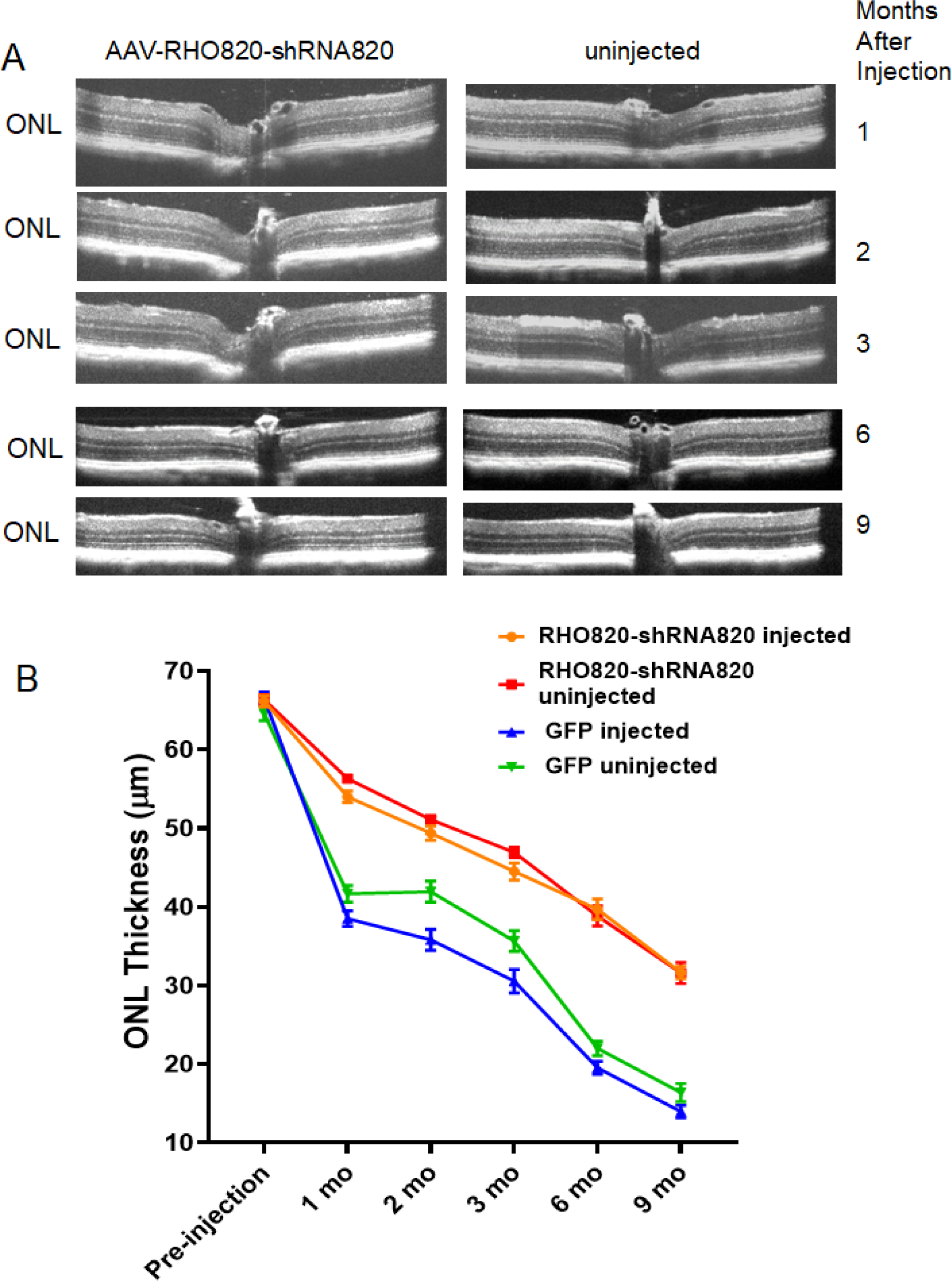
AAV-RHO820-shRNA820 treatment preserves retinal structure in both eyes of unilaterally treated mice. **A**. SD-OCT B-scans of mice treated in their right eyes with AAV-RHO820-shRNA820 and untreated in their left eyes at the times indicated. **B**. We measured ONL thickness as described in Fig. 3B at the indicated intervals. Differences between injected and uninjected eyes in each treatment group ( AAV-RHO820-shRNA820 or AAV-GFP) lacked statistical significance at all time-points, with the exception that the ONL in AAV-GFP injected eye were thinner at the 2 month time point than the uninjected eyes of the same mice (p<0.05; n=10-15 mice).

Electroretinography also indicated bilateral benefit from the unilateral injection of AAV- RHO820-shRNA820 (**Fig. 6**). There was no significant difference between the dark-adapted (scotopic) ERG amplitudes elicited between AAV-RHO820-shRNA820 treated eyes and uninjected eyes of the same mice, either for the a-wave amplitudes (**Fig. 6A**) or the b-wave amplitudes (**Fig. 6B**). In contrast, treatment with AAV-RHO820-shRNA820 conferred some benefit on cone function, as photopic ERG amplitudes were higher in treated eyes than in untreated contralateral eyes (Fig. 6C). The benefit to contralateral eyes was not an artifact of injection, since AAV-GFP treatment did not increase the ERG amplitudes of contralateral eyes, and both eyes of AAV-GFP treated mice exhibited a reduced response relative to the eyes of AAV-RHO820-shRNA820 treated mice.

**Fig. 6.**
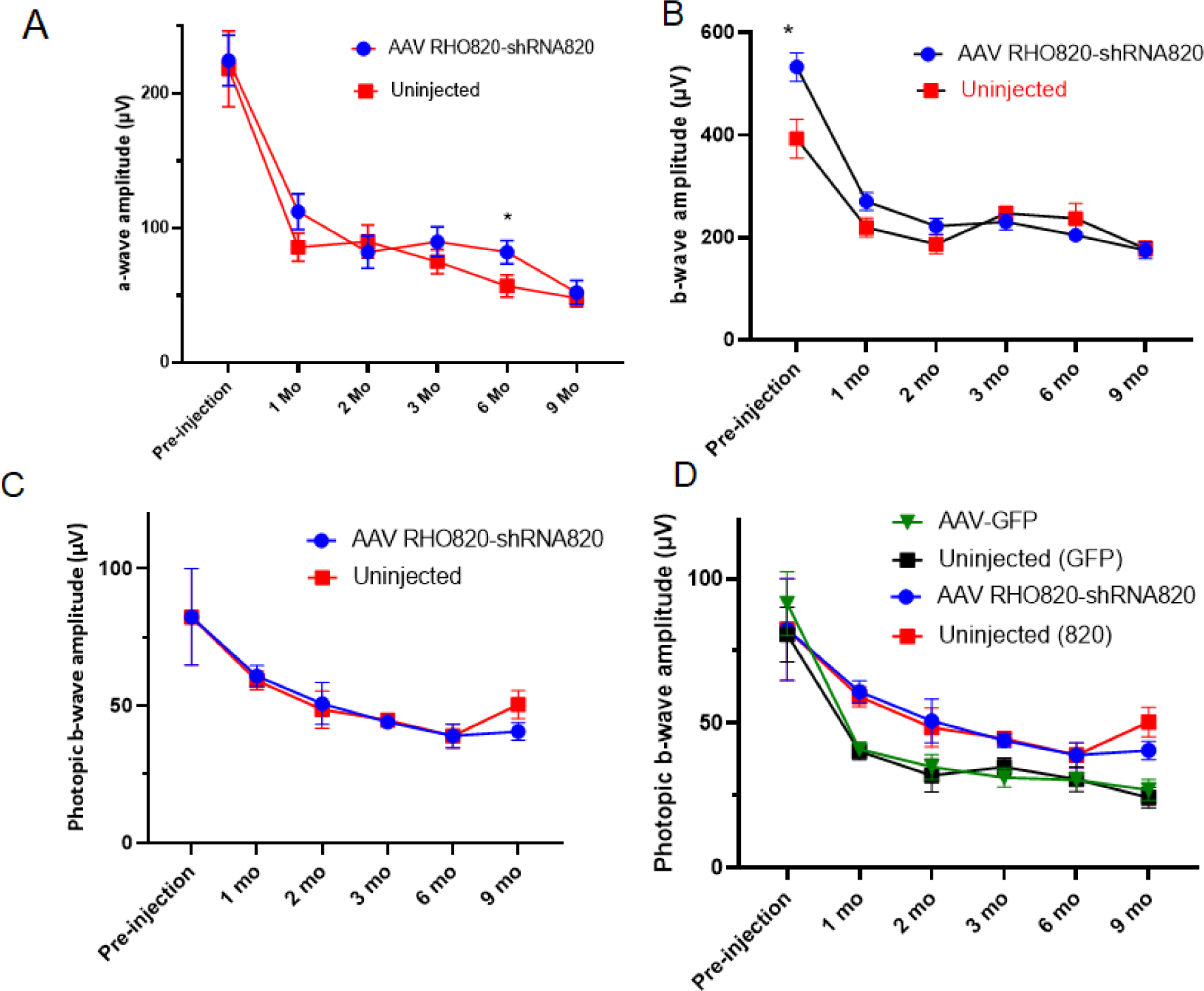
Treatment with AAV-RHO820-shRNA820 leads to improved retinal function in both eyes of unilaterally treated mice. **A**. dark-adapted a-wave amplitudes; **B.** dark-adapted b-wave amplitudes; **C.** light-adapted b-wave amplitudes. The data for AAV-RHO820-shRNA820 injected mice are the same as in Fig. 3A-C. **D.** For photopic amplitudes, differences between injected and uninjected eyes were not significant for eitherthe AAV-RHO820-shRNA injected mice or the AAV-GFP injected mice.

However, the difference was not statistically significant all time points (**Fig. 6D**).

### Is AAV transferred to the contralateral eyes?

To determine if the bilateral rescue that we observed could be attributed to the transfer of AAV from the injected eyes to the uninjected eyes, we first examined both eyes of the AAV-GFP injected mice by Western Blot of retinal extracts using an anti-GFP antibody. We had not observed direct GFP fluorescence by fluorescent fundoscopy (**Supplemental Fig. 1**) and no immunoreactive material was detected at a typical exposure of the immunoblot (**Fig. 7A**). However, an over-exposed blot revealed a small amount of GFP in the uninjected retinal samples (**Fig. 7B**). To attempt to quantify AAV transfer between eyes, we performed qPCR to estimate vector concentration in injected and uninjected eyes of mice treated with AAV-GFP. Our results (**Fig. 7C**) reveal that GFP mRNA was present in the uninjected eyes, but at a 10,000-fold lower level than in the eyes injected with AAV-GFP. We then sought to determine if AAV expressing *RHO820* was detectable in contralateral eyes. For this, we used DNA extracted from injected and uninjected eyes and primers from regions of the vector (hRHO promoter and SV40 intron) not included in the mature mRNA. We detected some viral DNA in the uninjected eyes, but at a level 3000-fold lower than injected eyes (**Fig. 7D**). To determine if *RHO820* mRNA could be detected in the uninjected eyes, we performed quantitative reverse transcription PCR using primers unique to RHO820 both within the coding region (RHO820-F, Table 1) and in the 3’- UTR (RHO820-R, Table 1). We detected expression of *RHO820* mRNA in uninjected eyes (n= 5), but the injected eyes expressed a 1300-fold higher level of RHO820 than the uninjected eyes (**Fig. 7E**).

**Fig. 7.**
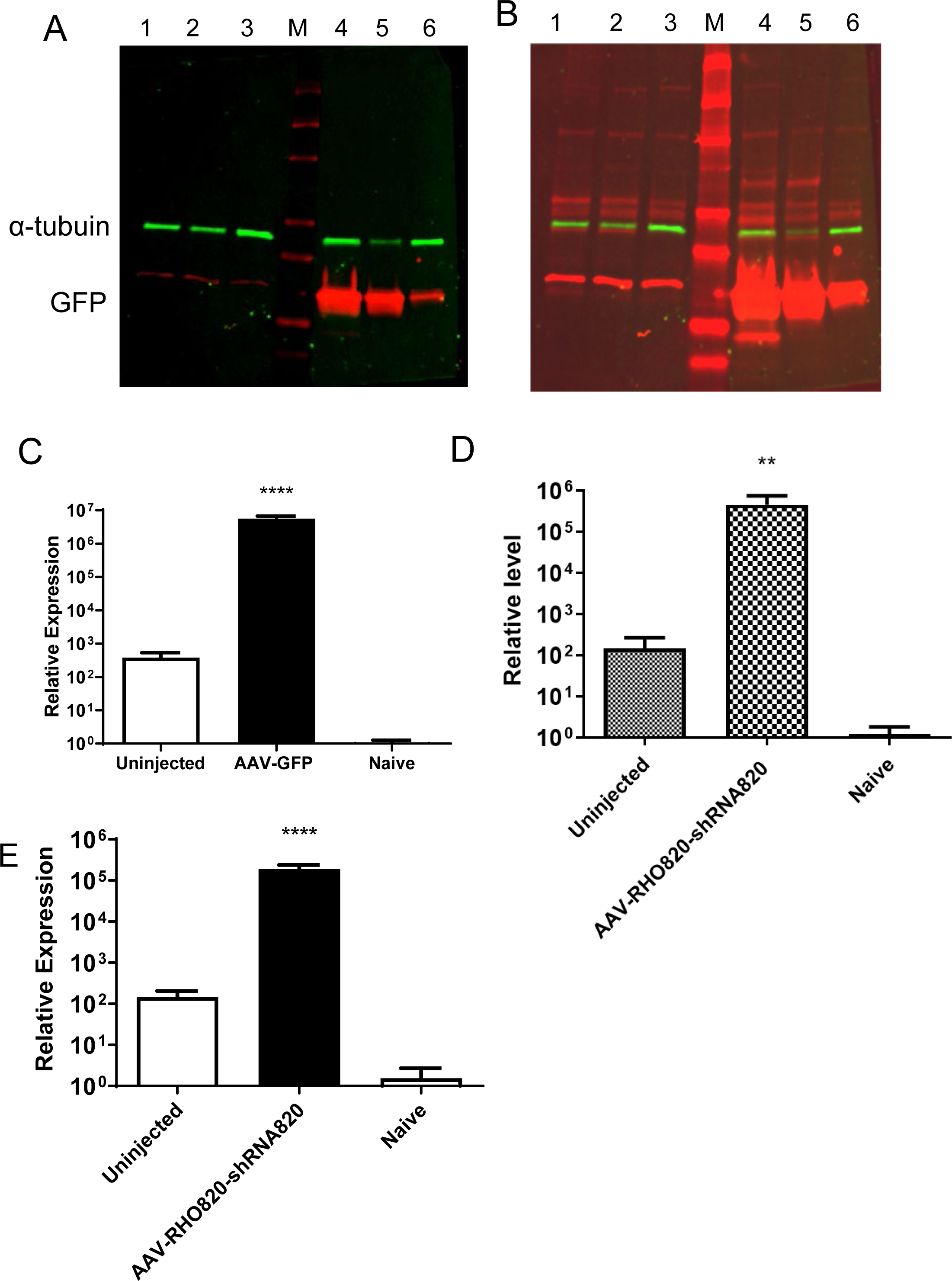
Detection of AAV mediated gene expression in the left eyes of mice injected with AAV-GFP or AAV-RHO820-shRNA820 only in their right eyes. **A**. Western blot of retinal protein extracted from uninjected left eyes (lanes 1-3) and AAV-GFP injected right eyes (lanes 4-6) of P23H *RHO* transgenic sacrificed one month post injection. M is a molecular weight marker. **B**. An over- exposure of the same blot as in (A). **C**. We extracted total RNA from right (injected) and left (uninjected) eyes of AAV-GFP injected mice, and amplified a GFP-specific sequence for quantitative RT-PCR. A low level of GFP was in the uninjected eyes, but no GFP RNA was detected in the eyes of untreated mice (naïve). *****=p<0.0001. (n=3) **D**. We used primers specific for the human opsin promoter and the SV40 intron (SD/SA) to detect vector DNA from both injected and uninjected eyes. No AAV-RHO820-shRNA820 DNA was detected in the eyes of naïve (untreated) mice. **E.** RNA was extracted from retinas of the AAV-RHO820-shRNA820 mice, and amplified with primers specific for *RHO820*. A low level of *RHO820* mRNA was detected in the left eyes of mice injected in the right eyes with AAV-RHO820-shRNA820. For graphs C-D, n=5.

## Discussion

Comparing AAV-RHO820-shRNA820 treated eyes with AAV-GFP treated eyes, we demonstrated that subretinal injection of this vector depleted human *RHO* mRNA and improved retinal structure and function in a mouse model of adRP caused by the P23H mutation in the human gene for rhodopsin. This result confirms that our approach is mutation-independent, in that our previous experiments with the same vector were performed in dogs with the T4R mutation.^7^ Both mutations lead to a similar biochemical phenotype: the opsin they encode tends to be retained in the endoplasmic reticulum and does not quickly reconstitute with 11-cis-retinal^13^. In addition, P23H *RHO* is associated with sector RP, in which the inferior hemisphere of the retina shows primary clinical signs of the disease^14^. This suggests that P23H, like the T4R mutation, causes light-induced degeneration of photoreceptors, albeit to a lesser extent. This sensitivity has been confirmed in animal models^15–17^. Since shRNA820 does not cleave the endogenous *Rho* mRNA (**Fig. 1**), the retinal protection we observed may be attributable to depletion of P23H *RHO* mRNA rather than augmentation with human RHO cDNA^18^.

In our previous experiments with AAV-Rho301-shRNA301 in P23H *RHO* transgenic mice, ERG a-wave and b-wave amplitudes were five-fold higher in the experimental eyes compared to eyes treated with AAV-GFP^2^. We speculate that the reduced functional protection with AAV- RHO820-shRNA820 is attributable to the expression of *human* rhodopsin from the human *RHO* promoter in the current vector instead of the murine rhodopsin from the proximal portion of the mouse *Rho* promoter used in our previous experiments. Even though the two proteins are 95% identical, the coding regions of the two mRNAs are only 88% identical, and differences in codon usage may lead to differences in expression. Another essential difference between the earlier experiments and those reported here is that in the current experiments we treated mice at one month of age in the current experiments, when retinal degeneration has already begun in this model. In our previous experiments, we injected mice just after their eyes opened, on postnatal day 14. Based on ERG responses, Cideciyan et al^19^ observed significant protection of the retinal function and structure in T4R *RHO* dogs, but those animals, too, were injected before the onset of retinal degeneration, which was stimulated by exposure to bright light.

While we did not anticipate bilateral protection from monocular injections, groups developing gene therapy for Leber Hereditary Optic Neuropathy (LHON) have reported improved visual function in the untreated eyes of patients injected unilaterally with AAV expressing a nuclear version of the mitochondrial ND4 gene^20–23^. Recently, Yu Wain-Man and colleagues reported the results of a long-term study of LHON gene therapy in 37 patients^23^. Of these, 29 exhibited significant improvement in best-corrected visual acuity in both eyes, though only one eye was treated with the AAV2-ND4 vector. Using cynomolgus monkeys, these authors report evidence of vector DNA in the retina and optic nerves of uninjected eyes. In a rat model, Yang et al. found evidence for the transfer of fluorogold nerve tracer from the injected eye to the anterior optic nerve of the untreated eye^24^. They did not find evidence of AAV transfer based on GFP fluorescence, however. It has been established that AAV can traffic to the brain via axonal transport along the optic tracts following intravitreal or sub-inner limiting membrane injection^25, 26^, and retrograde transport may explain the presence of vector in the optic nerves and retinas of injected macaques. That the extent of this transfer is sufficient to improve vision is both impressive and welcome.

In these experiments, we employed subretinal, rather than intravitreal, injection. While there are some reports of viral transfer to the brain following subretinal injections^27, 28^, transfer from one eye to the other has not been reported. Using the same vector used in these experiments, we saw no evidence for bilateral protection of dogs injected in one eye^7^. Indeed, we saw no benefit outside of the injection bleb (area of retinal detachment). It should be noted that subretinal injection of mice leads to pan-retinal transduction (**Supplemental Fig. 1**) whereas injection of dogs and humans leads to localized infection in the area of the retinal detachment. As mentioned above, gene therapy tests in rodents routinely use sham-injected fellow eyes as a control, with no evidence of structural or functional improvement in the sham-treated eyes^29^. Based on GFP immunoreactivity and DNA amplification, we found evidence of low-level transfer of AAV to the uninjected eyes (**Figs. 7 & 8**). Nevertheless, preservation of retinal structure, based on SD-OCT and histology, and of retinal function, based on dark-adapted ERG, was indistinguishable between treated and untreated eyes of the same mice (**Figs. 5 and 6**). qRT-PCR revealed a low level of expression of AAV-delivered *RHO820* mRNA in the uninjected eyes of mice treated with AAV-RHO820-shRNA820 in their contralateral eyes. In addition, total human *RHO* mRNA appeared to be diminished in the contralateral eyes of P23H transgenic mice injected with AAV-RHO820-shRNA820 (**Fig. 2A**). We conclude that this level of expression and knockdown was sufficient to reduce the rate of photoreceptor degeneration in the uninjected eyes. It will be interesting to see if bilateral protection is observed in human patients with RHO-related adRP treated by this method.

## Materials and Methods

### Experimental animals

All studies performed in adherence with the ARVO Statement for the Use of Animals in Ophthalmic and Vision Research and had prior approval of the University of Florida IACUC. P23H *RHO*::*Rho^+^/Rho^+^* mice were generated by breeding P23H *RHO* transgenic mice^30^ with C57Bl/6 BomTac mice (Taconic Biosciences, Rensselaer, NY, USA). The P23H *RHO* transgene was a 17.5 kb fragment of human genomic DNA described by Olsson and colleagues^31^, but the line was generated at the University of Florida transgenic core. It was bred to homogeneity on the C57Bl/6 BomTac background, and breeders were screened for the absence of mutations affecting retinal integrity (rd1, rd8)^32^ and for the presence of the M430 allele of *RPE65* characteristic of C57Bl/6 mice. Mice were routinely weaned at 21 days of age and injected with experimental viruses at 30 days.

### AAV vectors

We previously described AAV vector scAAV2/5-hOP-*RHO820*-H1-*shRNA820* (AAV-RHO820-shRNA820)^7^. It is a self-complementary AAV vector that encodes an human H1 promoter directing the synthesis of shRNA targeting human rhodopsin (Fig. 1D). It also contains an siRNA820 resistant *RHO* cDNA driven by the proximal human *RHO* promoter. The control vector was AAV expressing humanized GFP from the CBA (CMV enhancer, chicken beta-actin) promoter^33^. AAV2 vectors were packaged in AAV5 capsids and purified by the Ocular Gene Therapy Core using the methods of Zolotukhin et al.^34^

### Subretinal injections

We followed the methods of Timmers and colleagues.^35^ Mice were anesthetized with ketamine (95 mg/kg) and xylazine (5-10mg/kg) injected i.p in sterile saline. One drop of Tropi-phen (Phenylephrine HCI 2.5%–Tropicamide 1%) was used to induce mydriasis. The anesthetized animal was positioned on its side under a dissecting microscope. The cornea was punctured nasally approximately 0.5 to 1 mm anterior of the iris with a 28G hypodermic needle.

The puncture needle was removed carefully followed by the insertion of a blunt needle (33G) through the hole. With just the tip of the blunt needle entered into the anterior chamber, the syringe was turned such that the needle is pointed posteriorly. By gently sliding the needle in toward the posterior of the eye, the lens was pushed aside slightly to maneuver the needle-tip to the subretinal injection site. The animals were injected in one eye with one microliter of fluid, either with AAV- RHO820-shRNA820 or with AAV-GFP. Following injection, veterinary ophthalmic antibacterial ointment was applied to prevent drying of cornea and infection.

### Spectral Domain Optical coherence tomography (SD-OCT)

Spectral domain optical coherence tomography (SD-OCT) employed an Envisu instrument (Leica Microsystems, Buffalo Grove, IL, USA) as described by Ahmed et al.^36^ We averaged 25 B-scan images to reduce background and improve resolution. We made measurements of ONL thickness at four positions equidistant from the optical nerve head and averaged them to determine the mean ONL thickness for each eye.

### Electroretinography (ERG)

We performed ERG measurements as previously described^36^. We kept mice in a dark room overnight for scotopic measurements and then performed analysis under dim red light. We recorded responses from both eyes using an LKC UTAS Visual Electrodiagnostic System with a BigShot^TM^ full-field dome (LKC, Gaithersburg, MD). Scotopic ERGs were elicited with 10 msec flashes of white light at 0 dB (2.68 cds/m^2^), -10 dB (0.18 cds/m^2^), -20 dB (0.02 cds/m^2^) with appropriate delay between flashes. Five to 10 scans were averaged at each light intensity. To record light-adapted (photopic) ERG, we exposed mice to two minutes of white light. We used a flash of white light at 1.0 cd sec/m^2^ to elicit a cone response. The a-wave amplitudes were measured from baseline to the peak in the cornea-negative direction, and b-wave amplitudes were measured from cornea-negative peak to major cornea-positive peak.

### Histology

We harvested eyes for histology by removing the eyes by cutting the optic nerve. Eyes were stored overnight at −4°C in fixative (O-Fix; Leica, Buffalo Grove, ), followed by washing and storage in PBS. Eyes were embedded in paraffin, stained with hematoxylin and eosin, and sectioned by Histology Tech Services (Gainesville, FL, USA). We produced photomicrographs using a Leica DMi8 microscope.

### Knockdown Efficiency of human and mouse *RHO* mRNA

We seeded HEK293T cells in a 12-well plate and transfected them when the cells reached 70-90% confluence with 500ng of the CMV- RHO-turboGFP-BGH-PolyA plasmid expressing either human RHO or mouse RHO. Some wells also included 1μg of a self-complementary rAAV2 plasmid containing a control H1-shRNA cassette, an H1-shRNA820 cassette or an empty H1-shRNA cassette. Each co-transfection condition was performed in triplicate. After incubation, total protein was extracted and 15-20µg from each sample was separated on a 10% Mini-PROTEAN® TGX™ Precast Protein Gel (Biorad, Hercules, CA., USA) adjacent to the Li-COR Chameleon ladder (Li-Cor, Lincoln, NE, USA) and transferred to an iBlot PVDF Transfer Stack using Invitrogen’s iBlot system (Invitrogen, Carlsbad, CA, USA).

Membranes were blocked for 1 hour at room temperature with Odyssey blocking buffer (Li-Cor, Lincoln, NE, USA) and were then incubated with mouse anti-TurboGFP (1:2000; Origene, Rockville, MD, USA) and rabbit anti-β-tubulin (1:5000; Millipore, Burlington, MA, USA) in blocking buffer overnight at 4°C. Membranes were washed three times with 0.1% Tween-20 in PBS before being incubated with IRDye 800CW donkey-anti-rabbit and IRDye 680RD goat-anti-mouse (Both 1:5,000; Li-Cor, Lincoln, NE, USA) for 45 minutes at room temperature. Membranes were washed three times with 0.1% Tween20 in PBS and imaged with an Odyssey CLx system (Li-Cor, Lincoln, NE, USA). We used ImageJ software to measure band intensity (https://imagej.nih.gov/ij/). To measure the band intensity of the predicted monomeric form of RHO, we drew a box around the prominent band appearing at ∼65kDa. Band intensity of RHO was corrected for loading by measuring and dividing by the band intensity of β-tubulin. For experiments reported in Fig. 7, we normalized to α- tubulin. We report values as relative intensity, which was calculated as the corrected band intensity of each sample divided by the average corrected band intensity for the control shRNA condition.

Statistical significance was determined via one-way ANOVA followed by Tukey’s multiple comparisons test.

### Protein and nucleic acid extraction of mouse retinas

For protein extraction, we suspended mouse retinas in 100 ul of a lysis buffer consisting of 50 mM Tris-HCl (pH 8.0), 150 mM NaCl, 1% NP-40, and a protease inhibitor cocktail (Santa Cruz Biotechnology). We homogenized the tissue by sonication for 20 seconds (twice), followed by centrifugation at 12,000xg for 2 minutes. We collected the clarified supernatants, and estimated protein estimation using the Pierce 660 nm Protein Assay Reagent (Thermo-Fisher) followed by dilution to 1 µg/µl. For nucleic acid extraction from retinas, we homogenized retinas in Trizol reagent (Thermo-Fisher) We used the Direct-zol DNA/RNA kit from Zymo Research (Irvine, CA, USA) to isolate RNA and DNA, following the protocols recommended by the manufacturer.

### Quantitative RT-PCR from tissue sampes

Cells mechanically suspended and then washed with phosphate-buffered saline. We extracted total RNA using the RNeasy Mini Kit from QIAGEN, following the manufacturer’s instructions. We used 1µg of RNA to produces cDNA using the iScript cDNA synthesis kit from Bio-Rad (Hercules, CA). Primers used for RT-PCR are listed in Table 1.

The PCR reaction mixture contained cDNA template, SsoAdvanced Super mix containing SYBR green (Bio-Rad (Hercules, CA), and 3 μM gene-specific primers. After denaturation at 95° C for 2 min, we used 40 cycles of denaturation at 94° C for 15 seconds followed by annealing/elongation at 60° C for 30 seconds in a C1000 thermal cycler CFX96 real-time system (Bio-Rad). We normalized gene expression to levels of β-actin mRNA. We compared relative gene expression with untreated samples using the CFX96 software from Bio-Rad.

## Acknowledgements

This research was supported by a grant from the National Eye Institute to Michael Massengill (F30 EY027163-01), by an NIH SIFAR Grant (1S10OD028476), by an unrestricted grant from Research to Prevent Blindness and by the Shaler Richardson Professorship Endowment to Alfred Lewin.

## Supplementary Figures

**Supp. Fig. 1.** Fluorescence fundoscopy reveals 70% transduction of the central retina in injected eyes. We injected mice subretinally in the right eyes but left the contralateral eyes untreated. One month later, we made fluorescent fundoscopic images of both eyes. **A**. Injected eyes showed GFP fluorescence over 70% of the imaged region (central 30% of the retina). **B**. No GFP fluorescence was detected in the uninjected eyes.

**Supp. Fig. 2.** Photoreceptor inner and outer segments are protected in AAV-RHO820-shRNA820 injected eyes. Top: SD-OCT B-scan of an eye injected with AAV-RHO820-shRNA820, one month post-injection. Bottom: A similar scan of an eye injected with AAV-GFP, one month post-injection. IS/OS=reflective band corresponding to rod inner and outer segments.

**Supp. Fig. 3.** Retinal degeneration is similar in the injected and uninjected eyes of mice treated in one eye with AAV-GFP. SD-OCT B-scans made at 1, 2, 3, 6 and 9 months post-injection.

**Supp. Fig. 4.** Unilateral injection of AAV-RHO820-shRNA820 preserves retinal structure in both eyes, but unilateral injection of AAV-GFP does not. We injected AAV subretinally at one month of age and stained sections from mice sacrificed nine months later. **A.** The outer nuclear layer (ONL) is preserved in both eyes of animals injected with AAV-RHO820-shRNA820. **B**. Thinning of the ONL was observed in both eyes of AAV-GFP treated mice.

**Supplemental Table 1.**
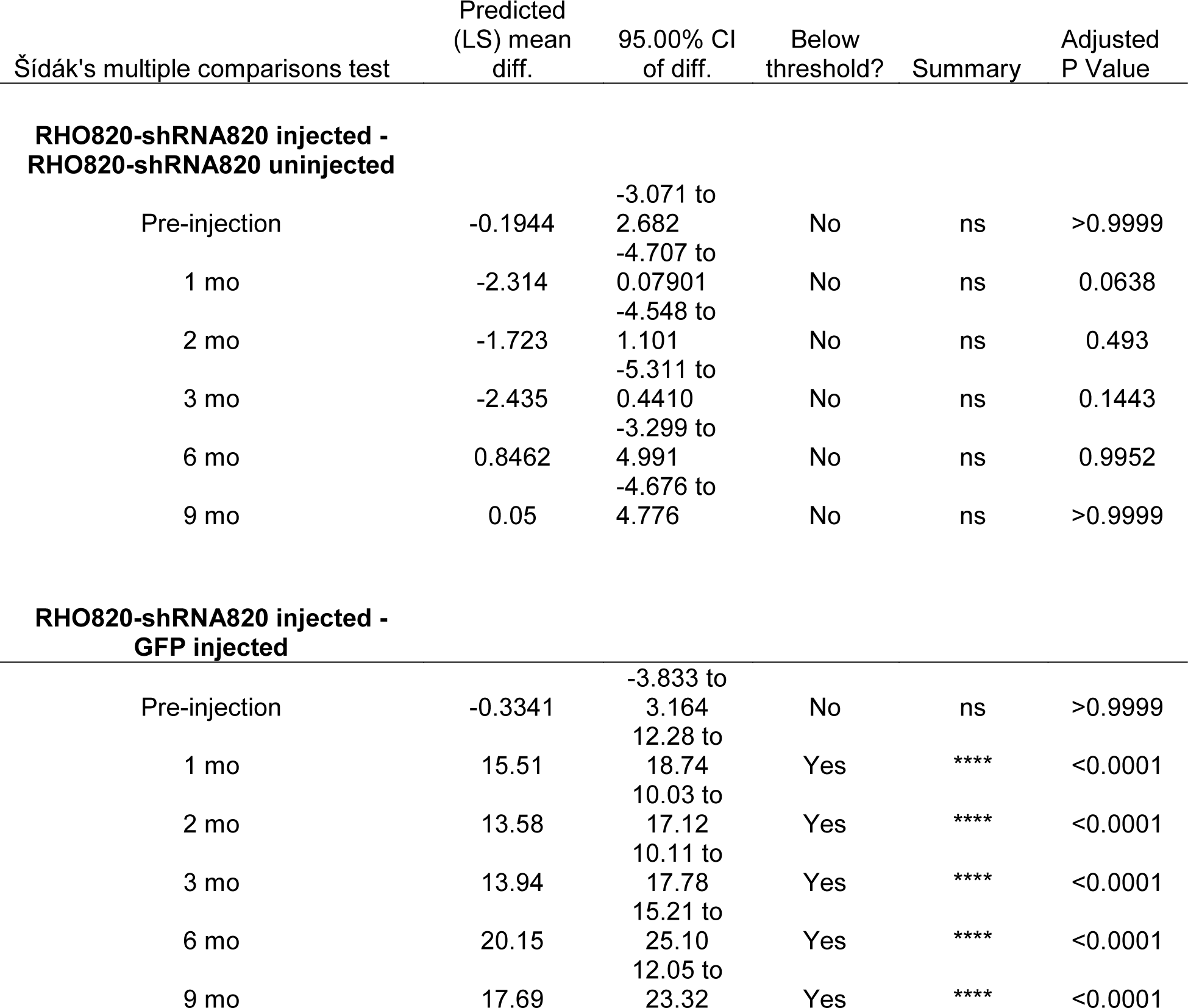

| Šídák's multiple comparisons test | Predicted (LS) mean diff. | 95.00% CI of diff. | Below threshold? | Summary | Adjusted P Value |
| --- | --- | --- | --- | --- | --- |
| **RHO820-shRNA820 injected - RHO820-shRNA820 uninjected** |  |  |  |  |  |
| Pre-injection | -0.1944 | -3.071 to 2.682 | No | ns | >0.9999 |
| 1 mo | -2.314 | -4.707 to 0.07901 | No | ns | 0.0638 |
| 2 mo | -1.723 | -4.548 to 1.101 | No | ns | 0.493 |
| 3 mo | -2.435 | -5.311 to 0.4410 | No | ns | 0.1443 |
| 6 mo | 0.8462 | -3.299 to 4.991 | No | ns | 0.9952 |
| 9 mo | 0.05 | -4.676 to 4.776 | No | ns | >0.9999 |
| **RHO820-shRNA820 injected - GFP injected** |  |  |  |  |  |
| Pre-injection | -0.3341 | -3.833 to 3.164 | No | ns | >0.9999 |
| 1 mo | 15.51 | 12.28 to 18.74 | Yes | **** | <0.0001 |
| 2 mo | 13.58 | 10.03 to 17.12 | Yes | **** | <0.0001 |
| 3 mo | 13.94 | 10.11 to 17.78 | Yes | **** | <0.0001 |
| 6 mo | 20.15 | 15.21 to 25.10 | Yes | **** | <0.0001 |
| 9 mo | 17.69 | 12.05 to 23.32 | Yes | **** | <0.0001 |

