## Supplementary figures and images for "Binocular Benefit Following Monocular Subretinal AAV Injection in a Mouse Model of Autosomal Dominant Retinitis Pigmentosa (adRP)"

### supplemental figures 1-4

Supplementary Fig. 1

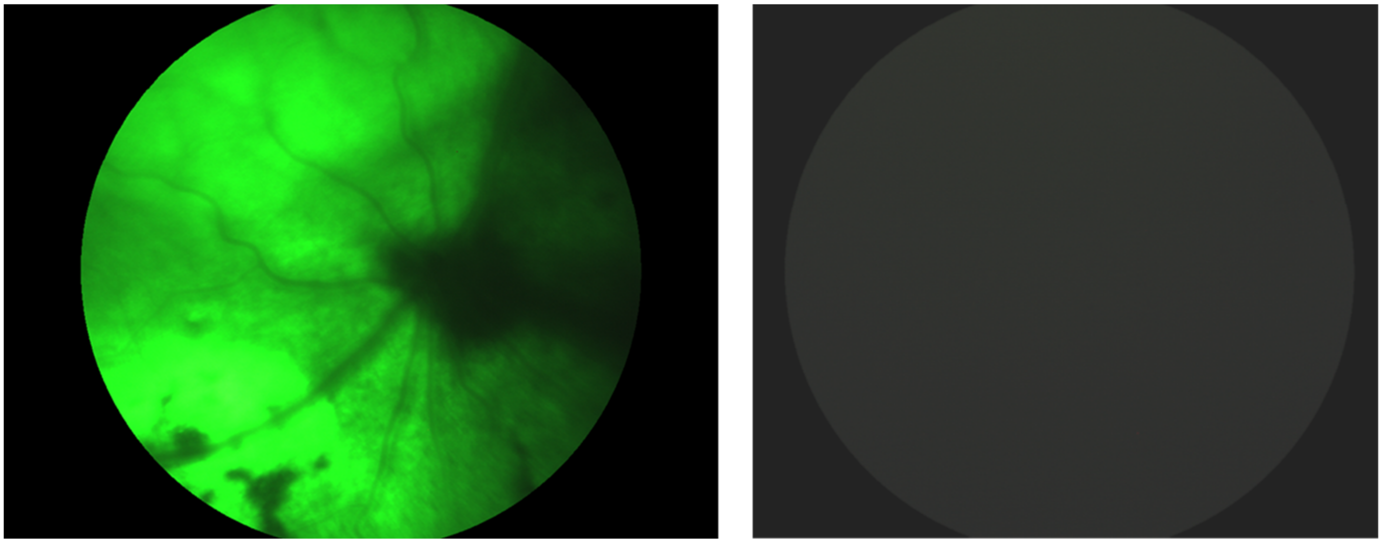

> 70% Coverage

Supplementary Fig. 2

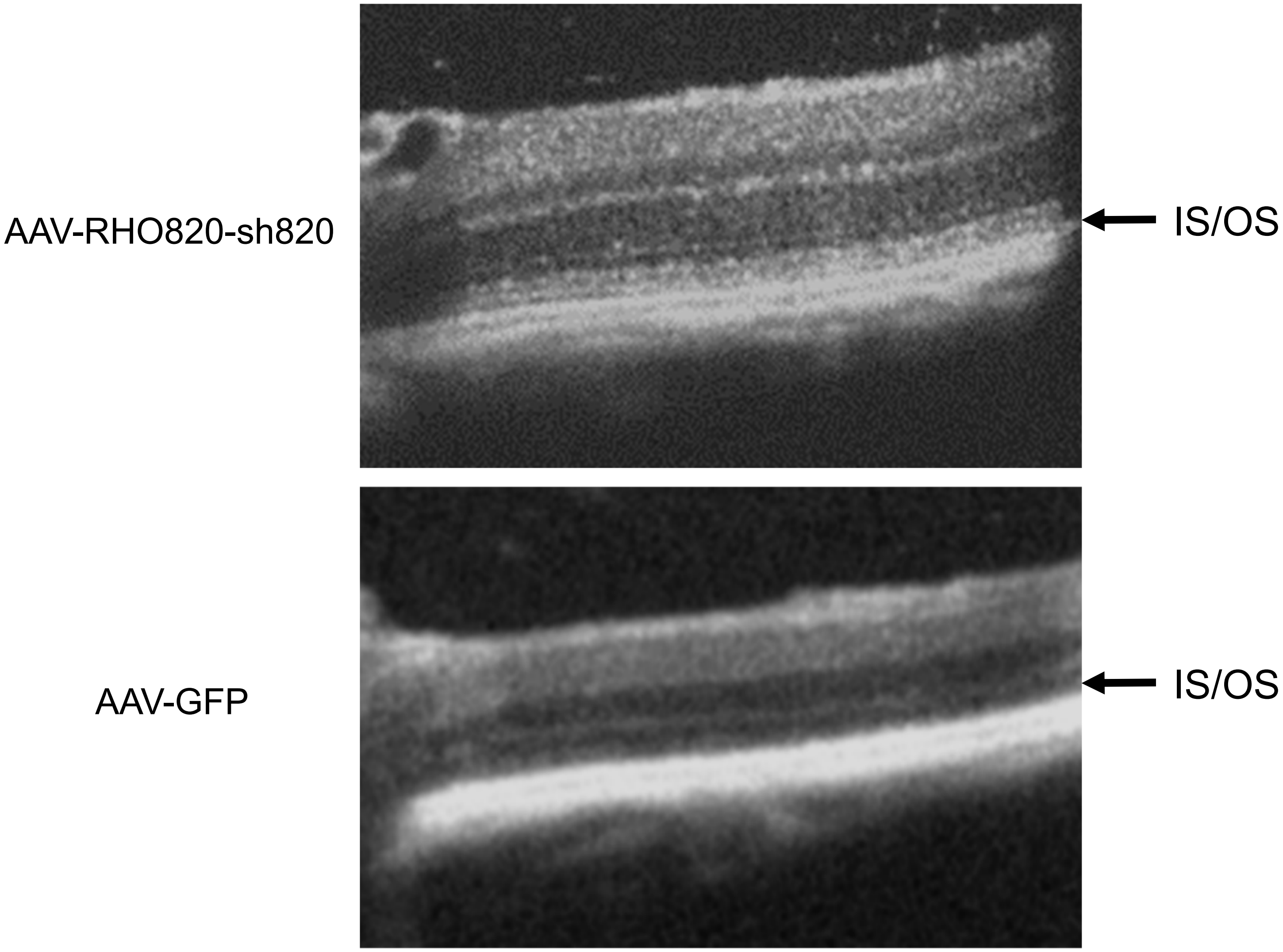

#### Supplementary Fig.3

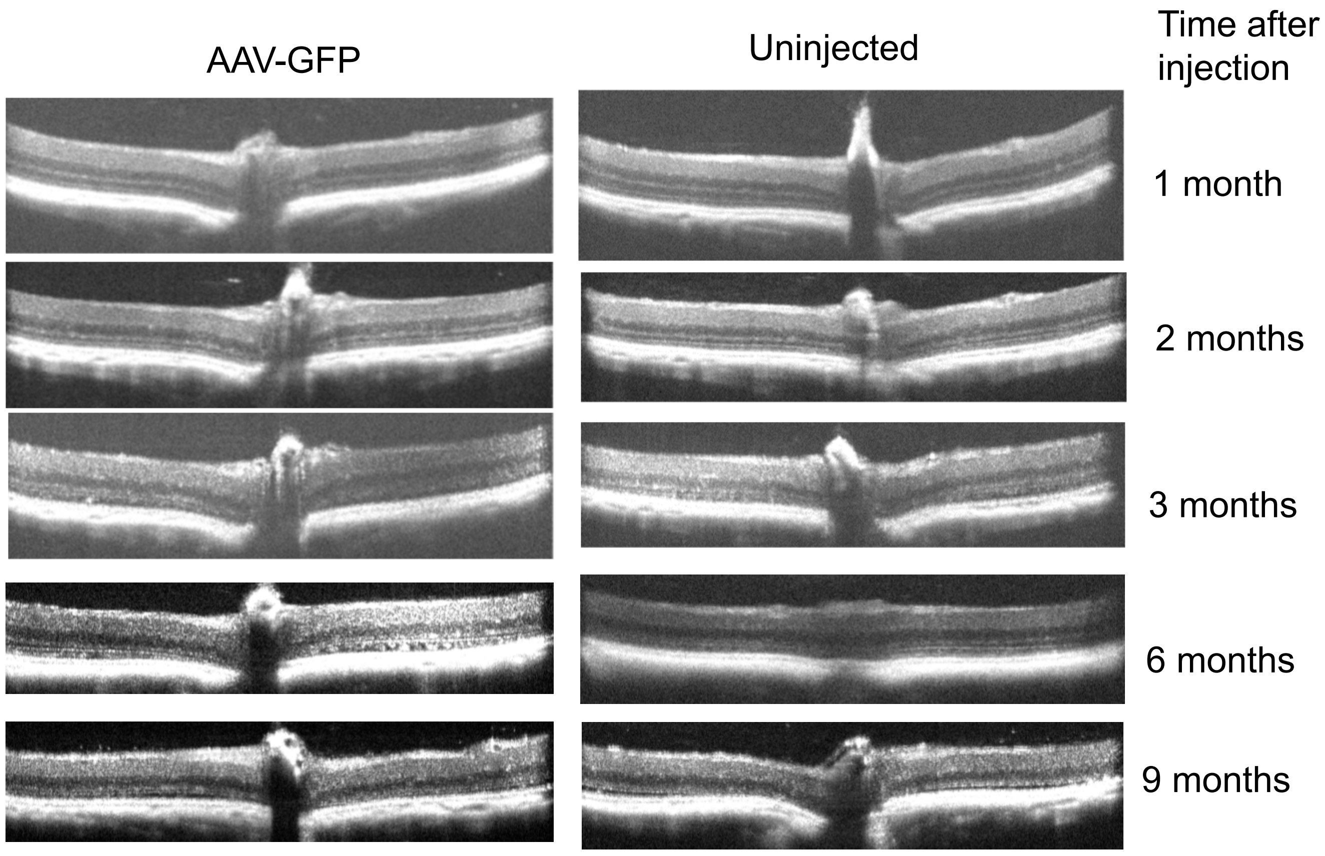

Supplemental Fig.4

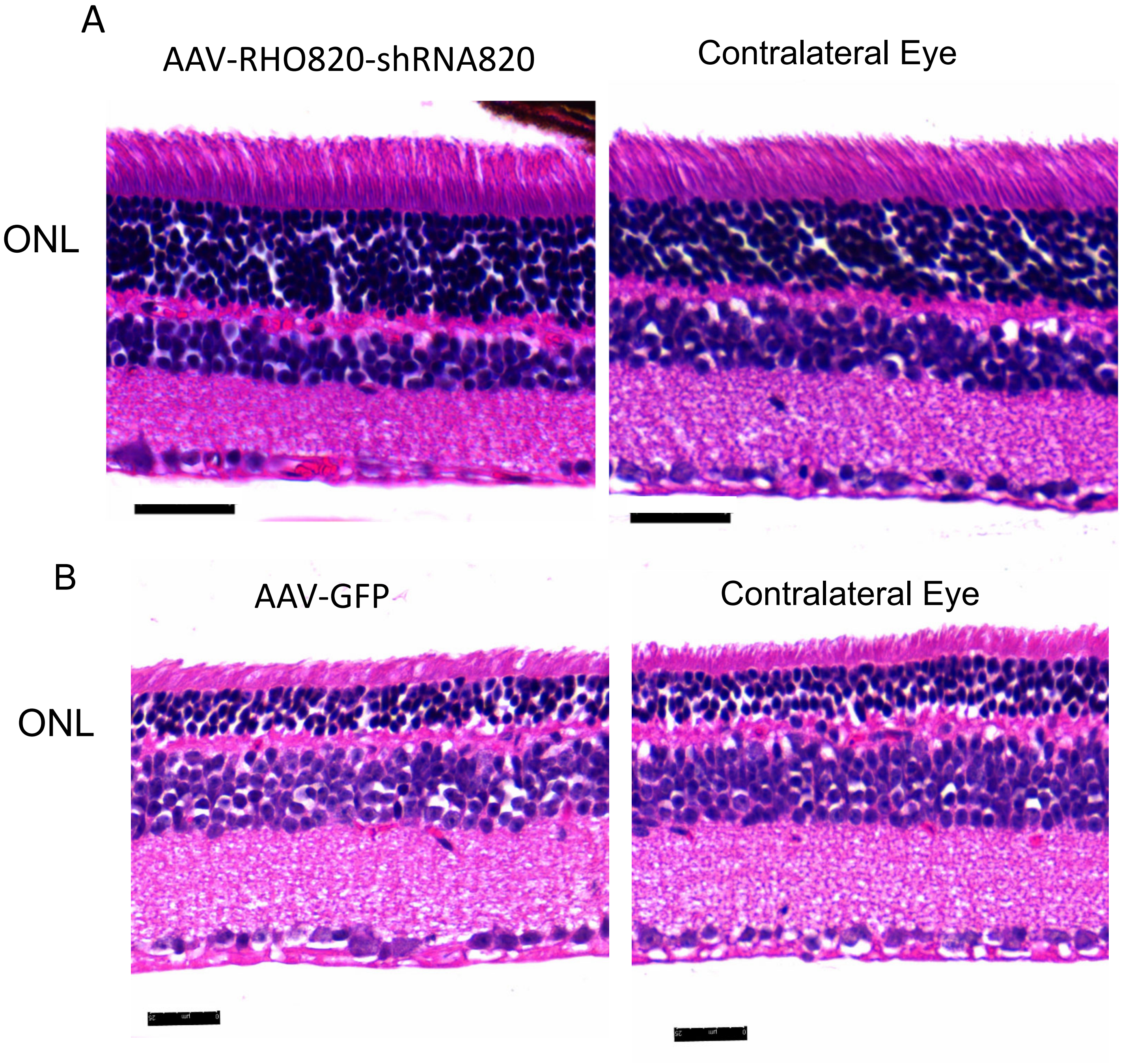
